# Organellar transcripts dominate the cellular mRNA pool across plants of varying ploidy levels

**DOI:** 10.1101/2022.03.12.484027

**Authors:** Evan S. Forsythe, Corrinne E. Grover, Emma R. Miller, Justin L. Conover, Mark A. Arick, M. Carolina F. Chavarro, Soraya C. M. Leal-Bertioli, Daniel G. Peterson, Joel Sharbrough, Jonathan F. Wendel, Daniel B. Sloan

**Author notes:** E.S.F and C.E.G. contributed equally to this work.

## Abstract

Mitochondrial and plastid functions depend on coordinated expression of proteins encoded by genomic compartments that have radical differences in copy number of organellar and nuclear genomes. In polyploids, doubling of the nuclear genome may add challenges to maintaining balanced expression of proteins involved in cytonuclear interactions. Here, we use ribo-depleted RNA-seq to analyze transcript abundance for nuclear and organellar genomes in leaf tissue from four different polyploid angiosperms and their close diploid relatives. We find that, even though plastid genomes contain <1% of the number of genes in the nuclear genome, they generate the majority (69.9–82.3%) of mRNA transcripts in the cell. Mitochondrial genes are responsible for a much smaller percentage (1.3–3.7%) of the leaf mRNA pool but still produce much higher transcript abundances per gene compared to nuclear genome. Nuclear genes encoding proteins that functionally interact with mitochondrial or plastid gene products exhibit mRNA expression levels that are consistently more than ten-fold lower than their organellar counterparts, indicating an extreme cytonuclear imbalance at the RNA level despite the predominance of equimolar interactions at the protein level. Nevertheless, interacting nuclear and organellar genes show strongly correlated transcript abundances across functional categories, suggesting that the observed mRNA stoichiometric imbalance does not preclude coordination of cytonuclear expression. Finally, we show that nuclear genome doubling does not alter the cytonuclear expression ratios observed in diploid relatives in consistent or systematic ways, indicating that successful polyploid plants are able to compensate for cytonuclear perturbations associated with nuclear genome doubling.

## INTRODUCTION

The endosymbiotic acquisition of mitochondria and plastids was instrumental in the early evolution and diversification of eukaryotes (1, 2). These organelles still retain their own genomes, but they are highly reduced in gene content owing in part to a combination of intracellular gene transfer to the nucleus and functional replacement via retargeting of existing eukaryotic genes (3–5). As a result, key functions in mitochondria and plastids, such as cellular respiration and photosynthesis, are now operated under divided genetic control, as they depend on intimate interactions between proteins encoded by different genomes within the cell (6, 7). There are now thousands of nuclear genes encoding proteins that are targeted to the mitochondria and plastids (8), which are known as N-mt and N-pt genes, respectively. Eukaryotes have evolved complex signaling mechanisms to coordinate organellar genome function with the expression, import, and turnover of N-mt and N-pt proteins (9–11). This regulatory task is complicated by the fact that nuclear, mitochondrial, and plastid genomes all have distinct transcription and translation systems (12).

Many of the protein subunits found in enzyme complexes that act in cellular respiration, photosynthesis, and other key organellar functions must assemble in equimolar ratios (Fig. 1). As such, disruption of the stoichiometric balance between N-mt/N-pt proteins and their cytoplasmic counterparts can have detrimental functional consequences (10, 13). Accordingly, eukaryotes must achieve this cytonuclear protein balance even though nuclear, mitochondrial, and plastid genomes differ wildly in copy number. For example, a typical diploid plant leaf cell contains two copies of the nuclear genome, dozens of copies of mitochondrial genome, and hundreds to thousands of copies of the plastid genome (14–17). Nuclear and organellar mRNA transcripts may exist in imbalanced ratios (18–21). For example, the mitochondrial OXPHOS genes have higher mRNA abundances than their N-mt counterparts even though the corresponding proteins are typically found in equimolar ratios within these complexes (18). Differences in nuclear vs. organellar transcript abundances appear to be related to intrinsic properties of the genomes themselves, as organellar genes that have been functionally relocated to the nucleus exhibit reduced transcript abundance (19).

**Figure 1.**
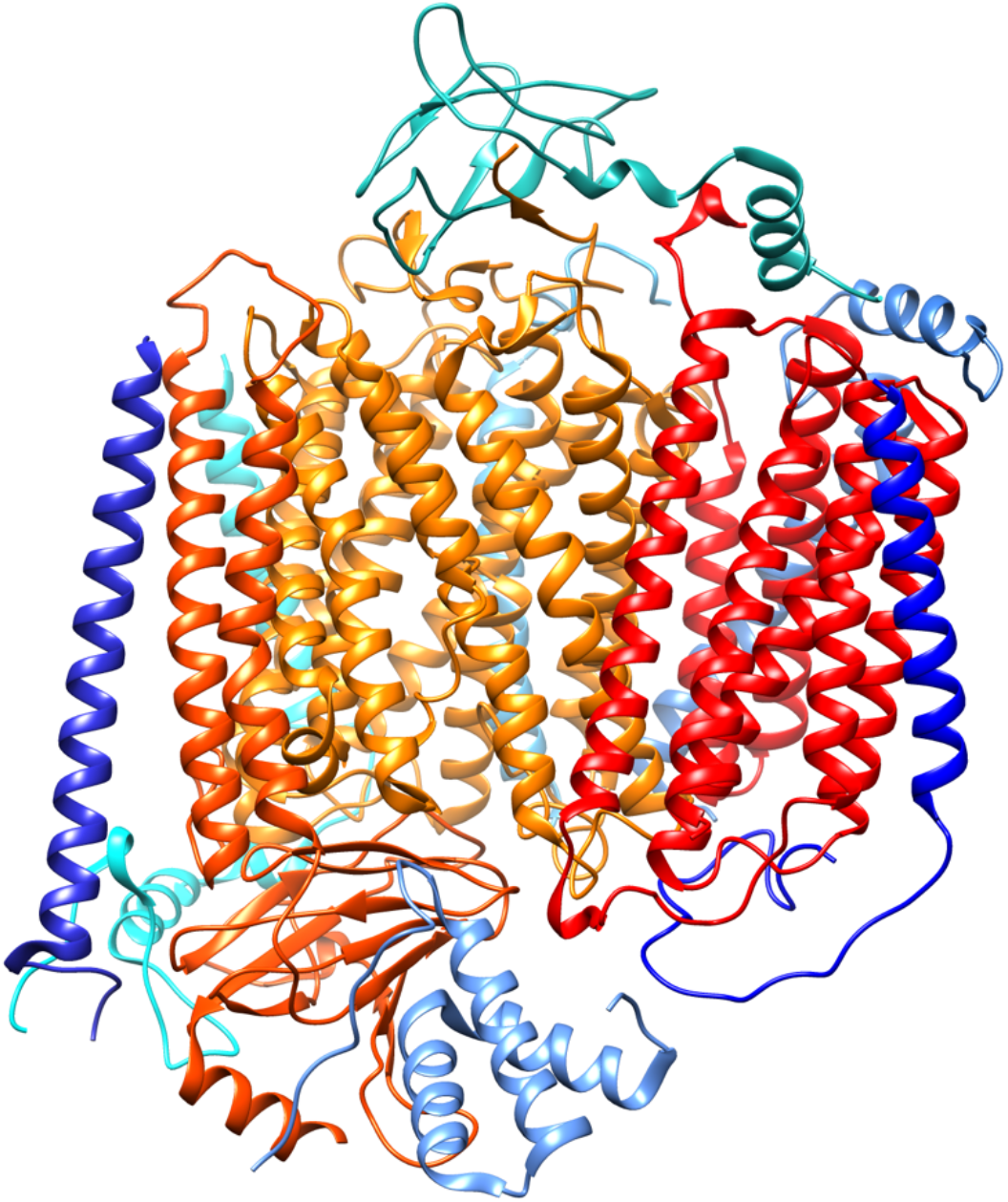
Structure of cytochrome c oxidase from the angiosperm *Vigna radiata* (70). Mitochondrial-encoded subunits Cox1, Cox2, and Cox3 are shown in shades of orange to red. Nuclear-encoded subunits are shown in shades of blue. All subunits assemble in an equimolar ratio with each present in a single copy within the complex. The image was generated with Chimera v1.15 (71), using Protein Data Bank accession 7JRO.

Doubling of the nuclear genome via polyploidy may further complicate the challenge of coordinating cytonuclear expression, as it has the potential to perturb the ratio of nuclear to organellar genome copies (22, 23). It appears that nuclear genome doubling elicits at least a partial compensatory increase in organellar genome copy number (15, 24–26), as polyploid cells often exhibit increased size and a larger number of organelles per cell (23). Whether the effects of nuclear genome doubling on cytonuclear stoichiometry are further stabilized by compensatory regulation of gene expression is less clear. A recent RNA-seq analysis of synthetic *Arabidopsis thaliana* autopolyploids indicated that they were able to preserve mRNA expression ratios for interacting nuclear and organellar genes (24); however, patterns differed between the mitochondria and plastids. The abundance of both plastid and N-pt mRNAs was reduced in a correlated fashion in polyploids relative to other nuclear genes and the diploid progenitor, whereas mitochondrial and N-mt mRNA levels were constant or even upregulated in polyploids. A more detailed breakdown of specific functional complexes and pathways within the organelles was not performed in that study, but another recent analysis used qPCR in diploid and polyploid *Leucanthemum* species to investigate mRNA abundance for plastid and N-pt genes encoding subunits of two specific photosynthetic enzyme complexes: photosystem II and RuBisCO (26). That analysis found that the two complexes differed with respect to whether cytonuclear mRNA balance could be restored in response to observed shifts at the genomic level in polyploids.

Given the variation observed in the foregoing studies, there is clearly a need for more global analyses of nuclear and organellar expression across multiple polyploid systems. A challenge in performing such analyses is to accurately quantify both nuclear and organellar transcripts in a total-cellular RNA pool. RNA-seq offers a means to quantify expression on a genome-wide basis. However, typical RNA-seq approaches in eukaryotes use polyA selection to exclude the highly abundant ribosomal RNAs (rRNAs) that would otherwise overwhelm sequencing datasets, rendering these approaches inappropriate for assessing organellar transcription. Whereas polyadenylation is a nearly universal feature of mature nuclear mRNAs, it is not ubiquitous in organellar mRNA processing. For example, in plant mitochondria and plastids, polyadenylation can occur, but it acts predominantly as a degradation signal (27, 28). As such, the polyA^+^ fraction of plant organellar mRNAs may tend to reflect transcript turnover rather than total expression (24). Ribosomal RNA depletion (ribo-depletion) is an alternative approach that can produce a less biased estimate than polyA selection for overall mRNA abundance between nuclear vs. organellar genomes, as well as expression differences among genes within these genomes. This method employs hybrid probes that specifically bind to and remove rRNAs without excluding other transcripts that lack a polyA tail (29).

In this study, we performed RNA-seq with ribo-depletion in four different angiosperm polyploids (*Arabidopsis suecica*, *Arachis hypogea*, *Chenopodium quinoa*, and *Gossypium hirsutum*). Each of these species is the product of allopolyploidization between two diploids, and high-quality genome sequences are available for all of them (30–33). We also analyzed diploid relatives that serve as models of the progenitors of each allopolyploid. The resulting data reveal a highly imbalanced cytonuclear stoichiometry in plant cells that is dominated by plastid and mitochondrial mRNAs. In addition, we find that these ratios are largely preserved between polyploids and related diploids, indicating that polyploid plants often compensate for cytonuclear perturbations associated with nuclear genome doubling.

## RESULTS

### Plastid transcripts represent the dominant fraction of the mRNA pool in plant leaf cells

We used ribo-depleted RNA-seq data and annotated nuclear, mitochondrial, and plastid protein-coding gene models to estimate mRNA abundance for each gene in terms of “transcripts per million” or TPM (34, 35). Even though nuclear genomes contain tens of thousands of protein-coding genes compared to only dozens in organellar genomes, we found that organellar transcripts dominated the mRNA pool in plant leaf cells (Figs. 2A., S1, and S2). In particular, the plastid genome accounted for the majority of total-cellular mRNA transcripts in leaf tissue from all 12 sampled species across four different angiosperm genera, with a mean of 76.2% and a range of 69.9% to 82.3% (Fig. 2A, Table S1). Mitochondrial mRNAs represented a substantially smaller percentage (mean of 2.2% and range of 1.3% to 3.7%; Table S1). Nevertheless, given the small number of genes in the mitochondrial genome, the average transcript abundance per gene is still much larger than for nuclear genes (Fig. 2B).

**Figure 2.**
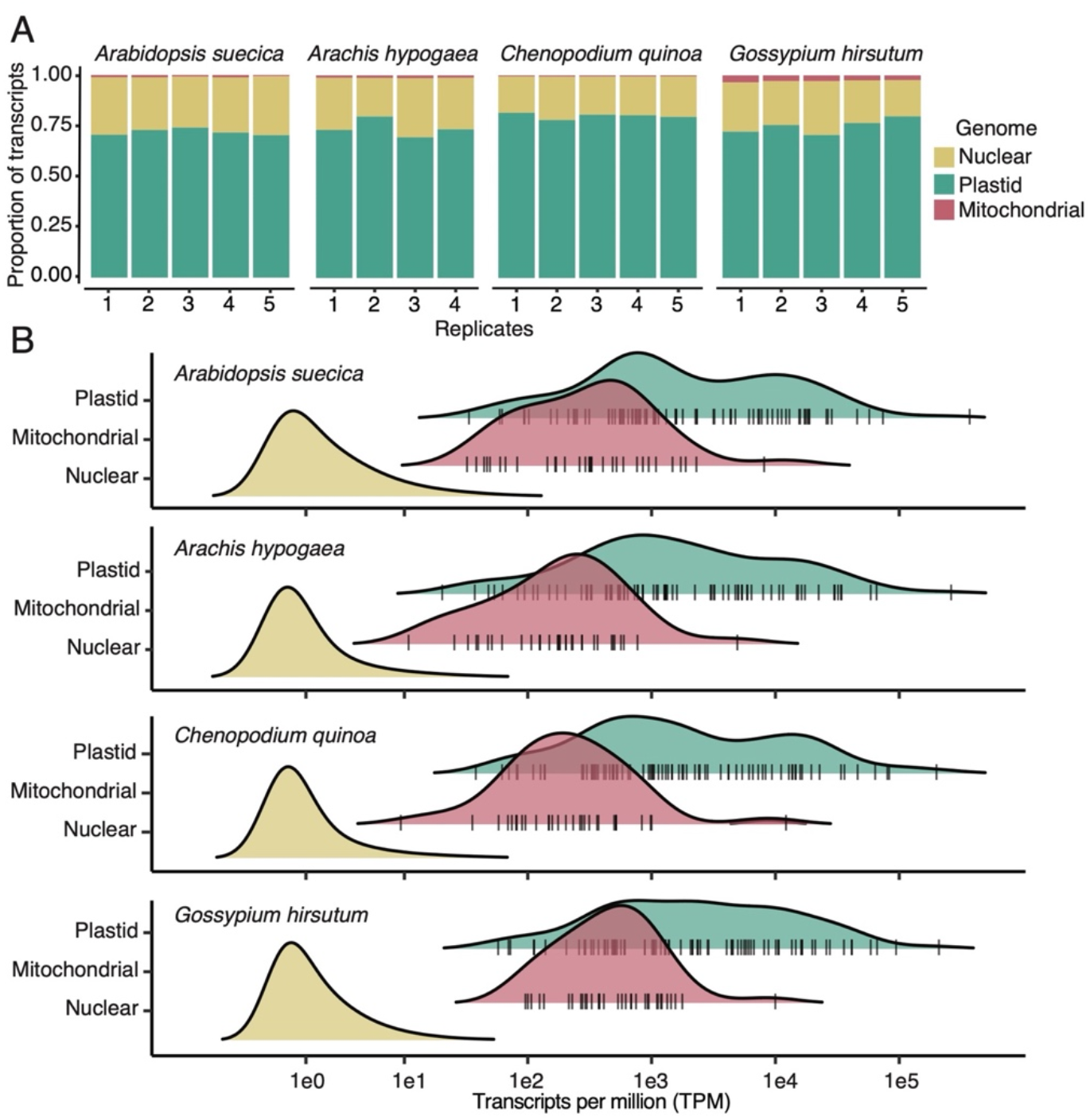
Predominance of organellar transcripts in the mRNA pool of plant leaf cells. A) Proportion of mRNA transcript by genomic compartment for each biological replicate in four allotetraploid species. B) Frequency distributions for transcript abundances by gene for each genomic compartment in each species. Vertical tick marks underlying the plastid and mitochondrial distributions indicate individual gene values. See Figs. S1 and S2 for corresponding data in diploids.

The abundance of mRNA transcripts varied dramatically across genes within the plastid and mitochondrial genomes, likely reflecting differences in rates of RNA polymerase binding and transcription (36). Photosynthetic genes were particularly dominant. In all species, the *psbA* gene, which encodes the D1 subunit of the photosystem II reaction center (37), exhibited the highest mRNA abundance, accounting for more than 25% of all mRNA transcripts in some samples (Table S2). The other most highly expressed plastid genes all encode subunits of key photosynthetic enzyme complexes (cytochrome b6f complex, photosystem I, photosystem II, and RuBisCO; Table S2). At the other extreme, the plastid genes with the lowest mRNA abundances generally encoded proteins involved in transcription (RpoB, RpoC1, and RpoC2) (38) or implicated in protein import (Ycf1 and Ycf2; Table S2) (39, 40), but these still produced higher mRNA levels than most nuclear genes (Fig. 2B). In mitochondrial genomes, the genes encoding subunits of the major oxidative phosphorylation (OXPHOS) enzyme complexes exhibited the highest mRNA abundances; *atp9* had the highest expression level in all species (Table S3). This gene encodes a subunit that is present in numerous copies that make up the membrane-bound c-ring within the FO portion of the mitochondrial ATP synthase complex (41). The lowest expression was observed for genes encoding proteins involved in cytochrome C biogenesis (CcmB, CcmC, CcmFc, CcmFn) (42), intron splicing (MatR) (43), and protein import (MttB; Table S3) (44).

Consistent with the predominance of photosynthesis-related transcripts in the plastid mRNA pool, the most abundant nuclear mRNAs generally encoded subunits in key photosynthetic enzyme complexes, particularly photosystem II and RuBisCO. However, as detailed below, the mRNA abundance for these nuclear-encoded subunits was far lower than for the corresponding plastid-encoded subunits that contribute to the same photosynthetic enzyme complexes.

### Extreme stoichiometric imbalance between mRNAs from organellar vs. interacting nuclear genes

For many of the major cytonuclear enzyme complexes in mitochondria and plastids, the constituent subunits must assemble in equimolar ratios. Examples include OXPHOS complexes I-IV, the ribosomes, the plastid acetyl CoA-carboxylase (ACCase), and key photosynthetic complexes such as RuBisCO, photosystem I, and photosystem II (8). Given this relationship at the level of protein subunits, the naïve or null expectation might be that there would be similar stoichiometric balance at the mRNA level. However, the massive transcript abundances observed for mitochondrial and plastid genes (Fig. 2) imply otherwise. To quantify nuclear expression for N-mt and N-pt genes involved in direct molecular interactions with organellar genomes or gene products, we analyzed major functional categories in the CyMIRA classification scheme (8). In order to have clear expectations for expression levels in diploids vs. polyploids, we restricted this analysis to nuclear genes found in “quartets”, meaning that they were present in two homoeologous copies in the focal allopolyploid and one copy in each of the two diploid progenitor models. To account for the fact that the gene copy number had doubled in the polyploids, we summed the transcript abundance values for the two homoeologs in polyploids and treated them as a single gene.

As predicted by the much higher TPM levels for organellar than nuclear genes (Fig. 2), overall mRNA abundance was heavily tilted towards mitochondrial and plastid genes, even when compared to their N-mt and N-pt counterparts involved in direct molecular and functional interactions. The mRNA abundance per gene was more than ten-fold higher for mitochondrial and plastid genes than for their interacting nuclear counterparts in all 12 species and all eight of the tested CyMIRA functional categories; in some cases, these cytonuclear ratios even exceeded 100-fold, including the mitochondrial ribosome and OXPHOS system in all *Gossypium* species and the plastid Clp complex in six different species (Figs. 3A, S3A, S4A).

**Figure 3.**
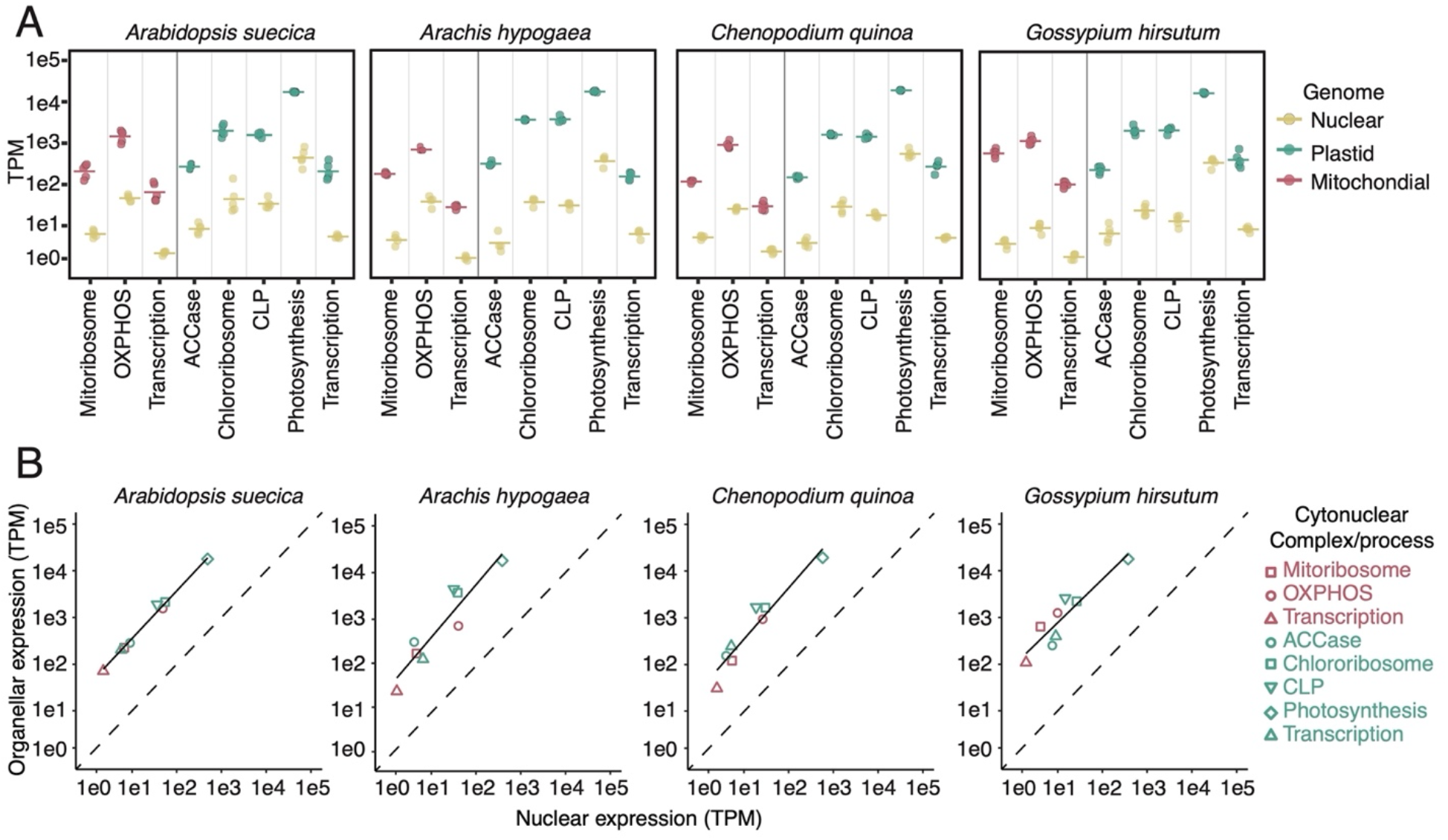
Consistent bias in cytonuclear stoichiometry for mRNA abundance. A) Transcript abundance for organellar genes and their interacting N-mt/N-pt counterparts broken down by functional categories according the CyMIRA classification scheme (8) in four allotetraploid species. Reported values are averaged across genes within a category. Points represent biological replicates, and horizontal lines indicate the mean across replicates. In all cases, organellar mRNA abundance (measured as transcripts per million or TPM) exceeds the corresponding nuclear mRNA abundance by at least ten-fold. B) Correlation between organellar and nuclear mRNA abundances across CyMIRA categories, with the solid line representing a best fit linear model, and the dashed line representing a 1:1 line. Despite the large imbalance towards organellar mRNAs, the cytonuclear ratio remains relatively consistent across functional categories expressed at very different levels, resulting in a strong positive correlation (*p* < 0.001 in all cases). For both sets of plots, the analysis of nuclear genes was limited to those found in homologous “quartets” (see Materials and Methods). Expression values for the two homoeologs in the polyploid reference genome were summed for this analysis. See Figs. S3 and S4 for corresponding data in diploids. See Figs. S3 and S4 for corresponding data in diploids.

### Nuclear and organellar expression levels are correlated across functional categories despite stoichiometric imbalance

One possible explanation for the massive cytonuclear mRNA imbalance is that mitochondrial and plastid genes are simply transcribed in excess and decoupled from the functional requirements for the corresponding gene products within the cell. To test this possibility, we compared average TPM values for organellar genes vs. interacting N-mt/N-pt genes across CyMIRA categories. In the absence of regulation at the transcriptional level in mitochondrial and plastid genomes, we would expect these values to be uncorrelated. To the contrary, this analysis revealed a strong positive relationship between organellar and nuclear mRNA abundance in each of the 12 species (*p* ≤ 0.001 for each of the 12 species; Figs. 3B, S3B, S4B). This finding represents clear evidence for cytonuclear coordination at the transcriptional level. The strong correlation across functional categories implies that expression levels in nuclear and organellar genomes both reflect cellular demands for these pathways despite the large observed stoichiometric imbalance that is biased towards mitochondrial and plastid mRNAs.

### Cytonuclear expression ratios remain stable across differing nuclear ploidy levels

The above analysis shows that the baseline expectation in plants should not be for balanced cytonuclear stoichiometry in mRNA abundance. Nonetheless, the strong correlation between interacting nuclear and organellar genes means that the observed cytonuclear ratio stays in a relatively narrow range across functional categories that differ greatly in absolute expression levels. This observation raises the question as to whether these ratios are robust to changes in nuclear gene copy number that accompany polyploidy events. To address this question, we first looked at whether the percentage of mitochondrial and plastid transcripts in the total-cellular RNA pool differed systematically between diploids and polyploids. All else being equal, doubling the nuclear genome copy number would lead to a proportional increase in nuclear mRNAs and concomitant decrease in the relative abundance of mitochondrial and plastid mRNAs. However, we found little evidence for a consistent reduction in the proportion of organellar transcripts in polyploids.

We found that the four polyploids each had total plastid mRNA abundance levels that were either intermediate between their two diploid progenitors (*Arabidopsis*, *Arachis*, and *Gossypium*) or higher than both of their diploid progenitors (*Chenopodium*), on average (Table S1). Therefore, none of the four systems exhibited the decline in plastid mRNA abundance relative to nuclear transcripts that might be expected to accompany nuclear genome doubling in the absence of any cytonuclear compensation, and there was only significant variation among species in plastid mRNA abundance in *Gossypium* (one-way ANOVA; *p* = 0.0031 in *Gossypium*; *p* ≥ 0.05 in the three other genera). For mitochondrial mRNA abundance relative to nuclear transcripts, there were significant difference among species within both *Arabidopsis* (*p* = 0.0042) and *Chenopodium* (*p* < 0.0001), but in each case the polyploid exhibited intermediate mitochondrial mRNA abundance compared to the two diploid progenitors on average (Table S1). For *Arachis* and *Gossypium*, the polyploid did exhibit lower relative abundance of mitochondrial mRNAs than both diploids, but these differences were not significant (*p* > 0.05 in both cases; Table S1).

Because the preceding analysis is based on total mRNA abundance in each organelle, it may be heavily influenced by individual genes with high expression levels such as *psbA* and *atp9* in the plastids and mitochondria, respectively. Therefore, we also compared mRNA abundance across species at the level of individual mitochondrial and plastid genes (represented as log10 TPM). In all cases, correlations in gene-specific expression levels between species were very strong (Figs. 4 and S5; Table S4). Patterns of variation between polyploids and diploids differed across genera, as well as between the mitochondria and plastids. In *Arabidopsis*, *post hoc* pairwise comparisons indicated that the polyploid *A. suecica* exhibited significantly higher gene-specific expression levels of plastid mRNAs compared to each of the two diploids (*p* < 0.0001), whereas it was intermediate between the two diploids for mitochondrial expression, with significantly higher values than *A. thaliana* (*p* < 0.0001) but significantly lower than *A. arenosa* (*p* < 0.0001). In the other three genera, the polyploid had lower organellar mRNA abundance values than both diploids (Table S4), although the difference between the polyploid *C. quinoa* and the diploid *C. suecicum* was not significant for mitochondrial transcripts (*p* = 0.06). Overall, the departures from a one-to-one relationship between diploids and polyploids were generally small and similar in magnitude to the differences between the two diploids (Fig. 4).

**Figure 4.**
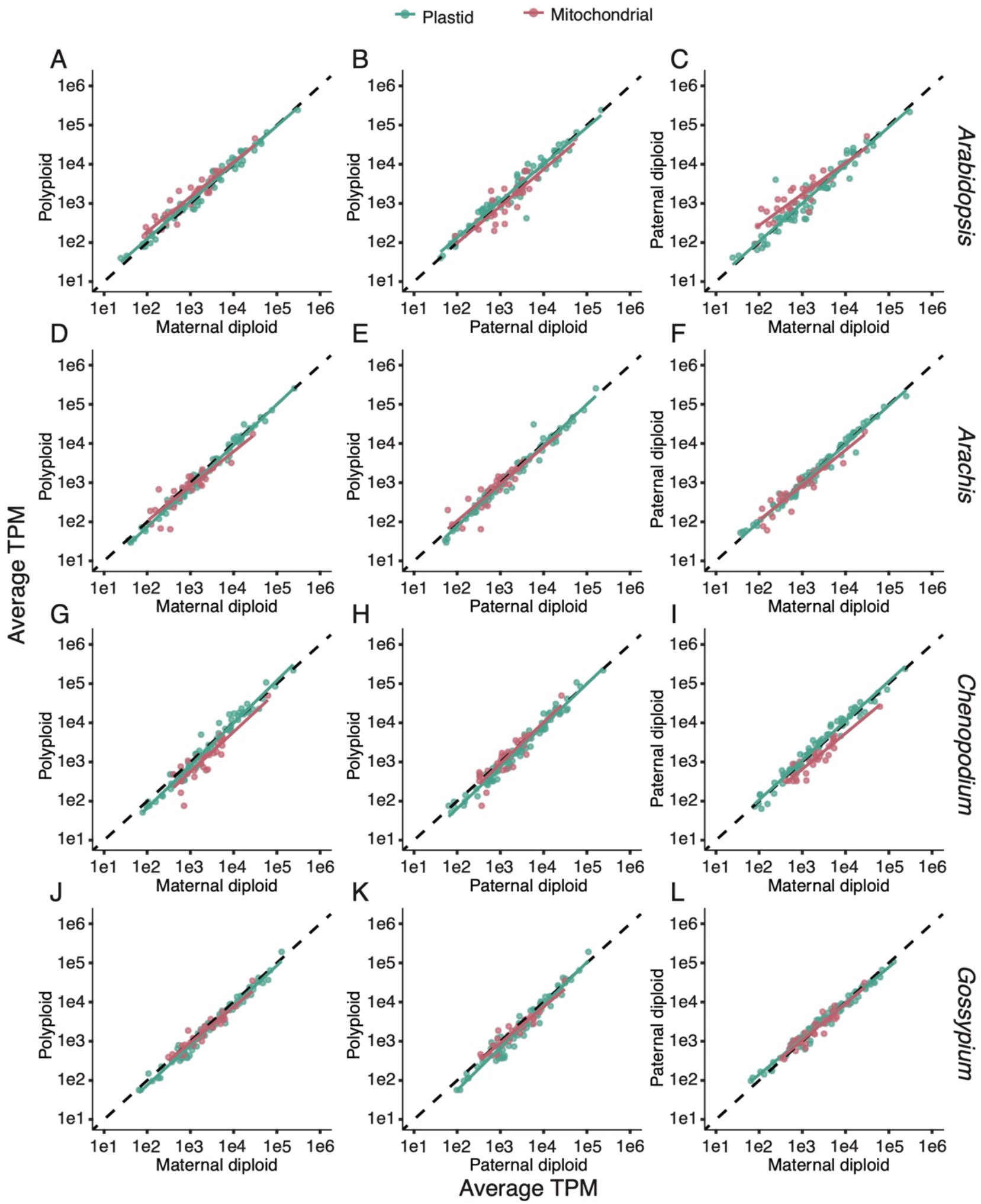
Comparisons between congeneric species, including diploids and polyploids, in relative abundance of organellar mRNAs, as measured by transcripts per million (TPM). The mitochondrial values are rescaled after removal of the plastid genes in each species, so they only reflect the relative abundance of mitochondrial and nuclear genes. Each point represents an average across biological replicates. The dashed line is the 1:1 line. Solid lines represent best-fit linear models for each organelle gene set separately. Each row corresponds to a different genus. Plots in the left column compare the polyploid against the maternal diploid. Plots in the center column compare the polyploid against the paternal diploid. Plots in the right column compare the paternal diploid against the maternal diploid. *R*^2^ values available in Table S4.

In addition to comparing mitochondrial and plastid mRNA abundance to total nuclear expression in polyploids vs. diploids, we also investigated how it specifically related to expression of N-mt and N-pt genes. Because of the functional relationships between organellar genes and N-mt/N- pt genes, we would expect any perturbation in these cytonuclear expression ratios in polyploids to have the most significant fitness consequences. We compared the ratio of expression levels for organellar genes to their interacting N-mt/N-pt genes between species. In most cases, these ratios exhibited a significant positive correlation across functional categories (Fig. 5). Similar to what we observed in analyzing total nuclear expression (Fig. 4), we did not find that polyploids from different genera showed consistent deviations in these cytonuclear expression levels (Fig. 5). For example, in *Arabidopsis*, the polyploid exhibited higher ratios of organellar to N-mt/N-pt transcript abundance compared to its maternal diploid progenitor (Fig. 5a), but the opposite was generally true in *Gossypium* (Fig. 5j). In addition, levels of N-mt/N-pt transcript abundance compared to other nuclear genes were similar between diploids and polyploids (Table S5). Overall, we do not find evidence that doubling the nuclear genome leads to consistent and long-term skewing of expression ratios towards nuclear genes.

**Figure 5.**
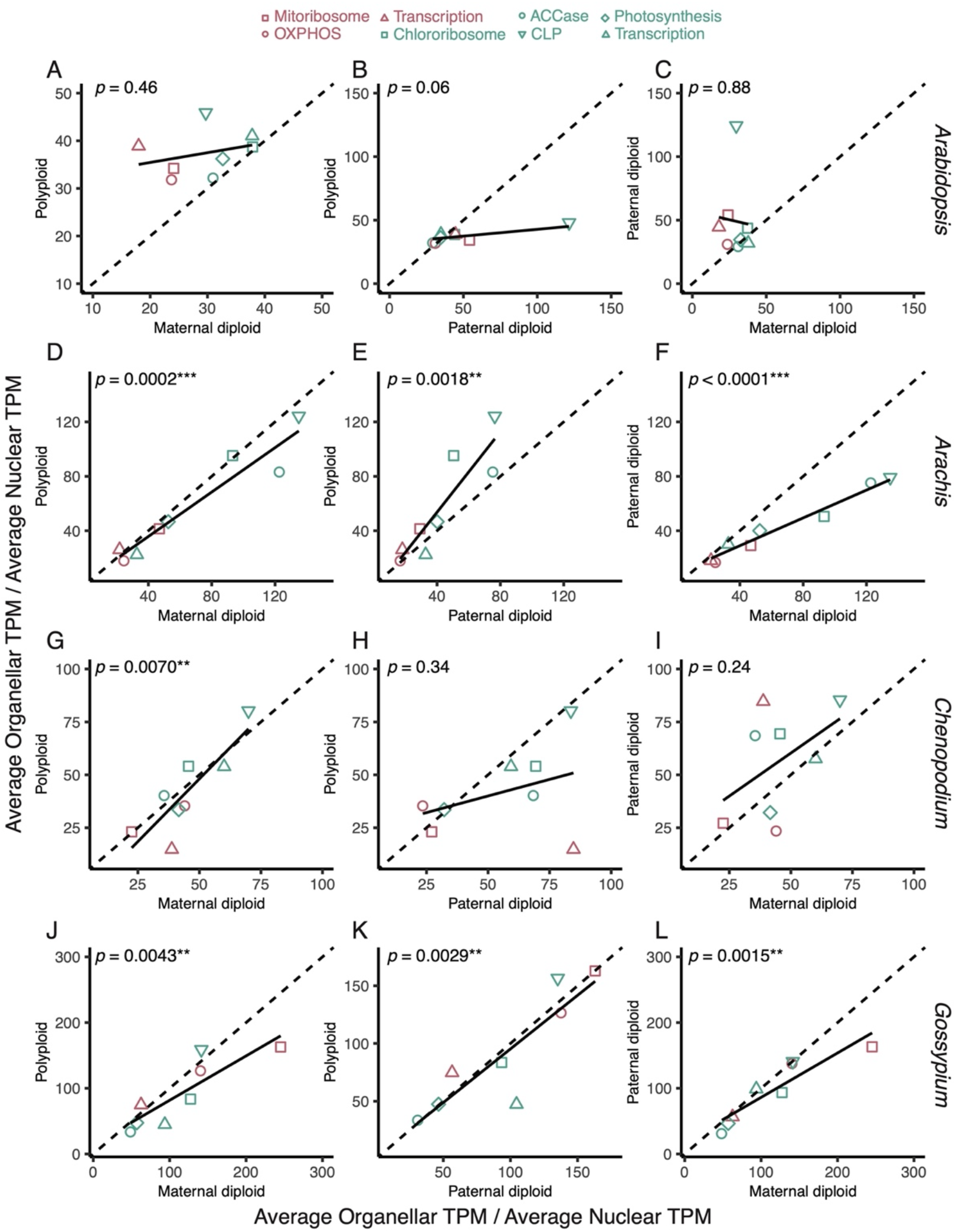
Comparison of cytonuclear stoichiometry in diploids vs. polyploids across functional classes. Scatter plots compare the ratio of organellar to nuclear mRNA abundance for individual CyMIRA categories between two congeneric species. Each point represents averages across biological replicates. The analysis of nuclear genes was limited to those found in homologous “quartets” (see Materials and Methods). Expression values for the two homoeologs in the polyploid reference genome were summed for this analysis. The dashed line is the 1:1 line, and the solid line represents a best-fit linear model. Each row corresponds to a different genus. Plots in the left column compare the polyploid against the maternal diploid. Plots in the center column compare the polyploid against the paternal diploid. Plots in the right column compare the paternal diploid against the maternal diploid.

## DISCUSSION

### The predominance of organellar transcripts in the plant leaf mRNA pool

One of the most striking patterns to emerge from our quantification of mRNAs across three different genomic compartments in plant cells is the sheer abundance of organellar transcripts (Fig. 2). Most plant RNA-seq studies use polyA selection, leading to a large underrepresentation of organellar transcripts (24, 28) and making direct comparisons between organellar and nuclear expression levels infeasible. In contrast to plants, animal mitochondrial mRNAs can undergo widespread polyadenylation (45). Accordingly, standard RNA-seq datasets have been used to quantify the abundance of mitochondrial vs. nuclear mRNAs in humans, and our findings parallel observations that the 13 protein-coding genes in the human mitochondrial genome can account for ~30-50% of all mRNA transcripts in tissues with high metabolic activity such as heart and brain (20, 21).

Given that chloroplasts can contain ~80% of the protein molecules in leaf mesophyll cells (46), it is not surprising that mRNAs with photosynthetic and other plastid functions predominate in plant leaf tissue. However, of the thousands of angiosperm genes that encode proteins in the chloroplast proteome, fewer than 80 (<5%) of them are in the plastid genome (47, 48). Therefore, the fact that plastid transcripts constitute the majority of the cellular mRNA pool can only be explained by the much lower expression levels for N-pt genes compared to their plastid counterparts. Previous studies have indicated that organellar transcripts can outnumber nuclear counterparts that encode proteins involved in direct functional interactions (18, 19). Our genome-wide analysis shows that this pattern extends across diverse functional categories with a wide range of expression levels (Fig. 3).

The imbalance in cytonuclear stoichiometry at the mRNA level contrasts with the equimolar ratios observed among protein subunits in many key cytonuclear enzyme complexes (e.g., Fig. 1). However, there are still multiple reasons that translational demands may be higher for mitochondrial and plastid-encoded proteins. First, even in cases where nuclear and organellar subunits co-assemble in a 1:1 ratio, they may be translated at different rates. For example, the fact that a remarkably large percentage of the entire mRNA pool in leaf tissue is dedicated to a single gene (*psbA*; Table S2) presumably reflects its rapid protein turnover and correspondingly high rates of translation. Even though the RuBisCO subunits are famous for being some of the most abundant proteins on Earth (49), the photosystem II D1 subunit encoded by *psbA* can be translated at higher rates in the light because of the frequent replacement of subunits damaged in photoinhibition (50).

Second, some N-mt and N-pt genes have been duplicated, affecting the amount of translation that would be expected per gene. For example, the plastid-encoded protein RbcL and the N-pt protein RbcS assemble in a 1:1 stoichiometric ratio to form the RuBisCO enzyme complex. However, there is only one copy of the gene encoding RbcL in the plastid genome, whereas RbcS can be produced by different members of a multicopy gene family in the nucleus (51, 52). Therefore, the amount of protein production per gene may naturally be lower for nuclear-encoded RbcS. Our analysis accounted for the recent gene duplications resulting from allopolyploidization by summing expression values for the corresponding homoeologs, but we did not adjust for older paralogous gene families in the nucleus. Therefore, such duplication events in the nucleus may contribute to imbalance measured on a per-gene basis (Fig. 3).

Third, the subunit composition of cytonuclear enzyme complexes deviates from equimolar stoichiometry in some cases. For example, the mitochondrial ATP synthase complex contains three copies of the Atp1 subunit and eight or more copies of the Atp9 subunit (53), both of which are mitochondrial-encoded in plants. Likewise, the sole plastid-encoded component (ClpP1) of the caseinolytic protease complex is present in three copies, whereas the corresponding N-pt subunits can be present in anywhere from one to three copies (54). Such cases are expected to increase demands for organellar expression. It also should be noted that, although all the functional categories that we analyzed are characterized by cytonuclear interactions at a molecular level, some of them do not involve multisubunit protein complexes with clearly defined stoichiometric relationships. For example, the mitochondrial-encoded MatR protein from the CyMIRA mitochondrial transcription and transcript maturation functional category does not form a complex with the N-mt proteins in that category, which include various proteins involved in mitochondrial transcription and post-transcriptional modifications (8).

Although each of the preceding factors are relevant to cytonuclear expression, it is unlikely that they can collectively explain the enormous imbalance between organellar and nuclear mRNAs, which consistently exhibited a difference of one to two orders of magnitude across diverse functional categories (Fig. 3a). Therefore, we are still left with the question as to why plants (and other eukaryotes) maintain such imbalanced cytonuclear expression at the mRNA level. One possibility is that organellar genomes are simply less reliant on transcription as a means of gene regulation (19). While it is apparent that standing levels of mRNAs are broadly tuned to the demand for specific organellar functions (Fig. 3b), it is less clear whether plant organellar genomes rely heavily on transcriptional regulation in response to short-term environmental changes. For example, upon shifting from dark to light conditions, there is a dramatic change in ribosome occupancy for *psbA* transcripts observed within minutes but minimal change in *psbA* transcript abundance (50). More generally, a recent analysis of high-light stress on chloroplast proteostasis found that, when rates of N-pt turnover increase, there is generally upregulation in mRNA abundance for the corresponding N-mt genes. In contrast, increased turnover of plastid-encoded proteins was rarely associated with an increase in the corresponding plastid mRNAs (55). These observations are consistent with the hypothesis that mitochondria and plastids are relatively lacking in precise, short-term transcriptional control and that the large standing population of organellar mRNAs offers a means for rapid regulatory responses at a post-transcriptional level.

One obvious difference between nuclear and organellar genomes is their copy number within the cell. Our estimates of transcript abundance “per gene” would be radically different if they were calculated per gene copy, given that plastid genomes are present in hundreds to thousands of copies per cell. Genome copy number, however, does not appear to be the sole determinant of the observed cytonuclear imbalance in mRNA abundance because mitochondrial-nuclear and plastid-nuclear ratios are similar to each other (Fig. 3) even though plastid genomes are present at much higher copy number than mitochondrial genomes in leaf tissue (14–17). Nonetheless, the ratios between organellar and nuclear mRNA abundances would be smaller and even imbalanced towards nuclear expression in some cases when tabulated per gene copy.

An intriguing feature of plant mitochondrial genetics is that some mitochondria contain only partial mitochondrial genome copies or no mitochondrial DNA at all (16). Intermittent mitochondrial fusion events may provide a means to restore complete genome content (56), but it is still unclear how these organelles retain function when they are lacking a full set of genes. Our observations raise the possibility that the abundance of mitochondrial mRNAs (Fig. 2b) may serve as a buffer to sustain protein expression through periods where the underlying genes are not available to be transcribed.

### The relative stability of cytonuclear mRNA stoichiometry across changes in nuclear ploidy

Previous work in these allopolyploids has found that homoeologs in both nuclear subgenomes are expressed at similar levels for most cytonuclear interaction functional categories (57). As such, doubling of the nuclear genome has the potential to alter the balance of cytonuclear gene expression, at least in principle (22). However, our analyses generally did not find consistent patterns of divergence in cytonuclear stoichiometry at the mRNA level between polyploids and related diploids, indicating that successful polyploids maintain cytonuclear stoichiometry at the transcriptional level following whole genome doubling. It is possible that potential perturbations in cytonuclear stoichiometry are alleviated by cellular and genomic responses even without changes to regulatory elements that control gene expression. In particular, elevated nuclear ploidy is associated with increases in cell size, number of organelles per cell, and the number of organelle genome copies per cell, resulting in at least partial compensation that stabilizes cytonuclear ratios at the DNA level (23). Notably, at least some of these responses can be observed immediately in lab-generated polyploids (15, 24).

Subsequent evolutionary responses in polyploid lineages may further fine-tune cytonuclear stoichiometry through regulatory changes in gene expression. For example, genome-wide changes in regulatory elements have been identified as a response to selection on stoichiometric balance following polyploidization (58). The long-term pattern of nuclear gene loss in polyploids also provides evidence for selection to maintain balanced cytonuclear expression, as N-mt and N-pt genes have been found to differ significantly in retention rates relative to the rest of the nuclear gene set (59–62). The use of natural allopolyploids in this study means that our polyploid-diploid comparisons reflect a combination of immediate developmental changes and longer-term evolution in response to nuclear genome doubling. Regardless of ploidy, divergence between any pair of species may lead to changes in the stoichiometry of cytonuclear gene expression, which is illustrated by the fact that observed differences between pairs of diploids were often comparable in magnitude to differences between polyploids and diploids in our dataset (Figs. 4 and 5; Tables S4 and S6). This variation highlights the importance of replicated sampling across multiple independent diploid-polyploid systems.

Although the larger number of organellar genome copies per cell in polyploids (15, 24–26) suggests that an increase in total organellar mRNA abundance contributes to the stability of cytonuclear expression ratios, it remains an open question as to whether there is also some suppression of nuclear expression after genome doubling in these systems. Therefore, an important area for future research is to characterize absolute levels of organellar and nuclear gene expression across different ploidy levels. Standard RNA-seq only measures relative gene expression, but it is possible to use RNA-seq to estimate changes in absolute expression levels per cell by quantifying the number of cells used in RNA extractions and applying spike-in controls during the sequencing process (63, 64). Given the increased cell size in polyploids, total transcriptome and proteome sizes are expected to scale allometrically with ploidy (65, 66), but how these scaling relationships affect many key features of cytonuclear interactions have yet to be explored in polyploid plants.

## MATERIALS AND METHODS

### Generation of RNA-seq data and mapping to reference genomes

The sampling of diploids and polyploids from *Arabidopsis*, *Arachis*, *Chenopodium*, and *Gossypium* is described elsewhere, including a summary of the history of allopolyploidization in each lineage, plant growth conditions, RNA extraction, sequencing, and read mapping (57). Briefly, RNA was extracted from the allopolyploid and its model diploid progenitors, ribo-depleted, and then sequenced as PE150 on a NovaSeq 6000 S4 flow cell. Reads from all three species within each genus were mapped to the annotated protein-coding sequences from the polyploid nuclear genome and from the organellar genomes of a representative member of the genus using Kallisto (67), as described elsewhere (57). We analyzed five biological replicates for each species with the exceptions of *Arachis hypogaea* and *C. suecicum*. For those species, one of the five replicates was excluded either because a preliminary clustering analysis failed to group the sample with other replicates from that species, raising doubts about sample identity (*C. suecicum*), or because of poor library quality and a low read mapping rate (*Arachis hypogaea*). For *Arabidopsis arenosa*, a total of seven biological replicates were available; however, we only included the first five to balance the sampling of the other species.

### Estimation of gene expression levels

Gene-specific TPM values for each sample were generated with Kallisto by using *kallisto quant*. Annotation quality has the potential to affect cytonuclear expression ratios in at least two significant ways. First, nuclear genomes contain numerous insertions of mitochondrial and plastid sequences (known as numts and nupts, respectively; (68)). In the case of recent insertions, true mitochondrial and plastid reads can map equally well to numt/nupt pseudogenes or to the actual organellar genomes, resulting in an underestimate of organellar expression. These cases were accounted for prior to read mapping by excluding numts and nupts that were annotated as protein-coding genes from each nuclear reference (57). Second, because we employed a ribo-depletion strategy rather than polyA selection, sequencing produced large quantities of reads from non-coding RNAs (ncRNAs) that lack polyA tails and would be excluded from typical mRNA-seq libraries (69), such as signal recognition particle RNAs and U1 and U2 spliceosomal RNAs. In some cases, the nuclear references contained annotations of short protein-coding genes corresponding to these ncRNAs whose high expression and short length led to extremely high TPM values. Because these high TPM ncRNAs could distort our estimates of nuclear mRNA expression, we manually screened all loci with values of ≥1000 TPM to identify protein-coding annotations that corresponded to highly expressed ncRNAs. These loci were retained in the reference for initial mapping to avoid mismapping of reads to other loci, but we subsequently excluded them and rescaled TPM estimates for the rest of the reference gene set so that the values summed to one million. Similarly, for certain analyses, we wanted to compare the relative abundance of nuclear mRNAs to plastid mRNAs by themselves or to mitochondrial mRNAs by themselves without the confounding effects of expression variation in the other organellar genome, so we excluded the relevant gene set and rescaled TPM values accordingly.

### Identification of homoeologous “quartets” and assignment of CyMIRA functional categories

For direct comparisons of nuclear gene expression between polyploids and related diploids, we used the set of previously identified gene “quartets” that were present as two homoeologous copies in an allopolyploid and a single copy in each of the corresponding diploid relatives (57). Because read mapping for all species was performed against the polyploid reference genome, we summed TPM values for the two homoeologs in each quartet for subsequent comparisons of expression. By summing these values, we were able to account for any mismapping of diploid reads to the other subgenome due to sequence similarity between homoeologs and to standardize for the difference in gene copy number between diploids and polyploids. For cross-species comparisons (Figs. 5 and S5), we excluded nuclear genes not assigned to quartets (57) and rescaled TPM estimates for the remaining gene sets accordingly. For consistency across analyses investigating cytonuclear expression ratios across CyMIRA categories, we also used this rescaled dataset for the analyses in Figs. 3, S3, and S4.

To assign nuclear genes to functional categories based on predicted targeting to mitochondria and plastids, we used the CyMIRA classification scheme (8) and a previously defined pipeline (62) (https://github.com/jsharbrough/CyMIRA_gene_classification). This classification scheme further assigns genes involved in direct molecular interactions with organellar transcripts or gene products to specific functional categories or enzymatic complexes (Figs. 3 and 5). Organellar genes were manually curated and assigned to the appropriate CyMIRA functional categories (Tables S2 and S3).

### Statistical analyses

Statistical analyses were carried out in R v3.5.0 or v4.0.5. Simple linear models (Figs. 3 and 5) and one-way ANOVAs were implemented with the lm and aov functions, respectively. For analysis of gene-specific expression levels across species, we used a mixed-model approach with species and gene as fixed effects and individual plant (replicate) as a random effect. This model was fit with the lmer function (lme4 package) and evaluated with the Anova function (car package) using type III sums of squares. *Post-hoc* comparisons between species pairs were conducted with the emmeans function, using a Holm procedure for the Bonferroni correction to account for multiple pairwise comparisons. These gene-specific models were applied after exclusion of genes from the other organellar genome and rescaling of TPM values as described above. Data visualizations were generated with the ggplot2 and ggridges packages. R code used to perform statistical analyses is available via https://github.com/EvanForsythe/Cytonuclear_RNAseq.

## ACKNOWLEDGEMENTS

This work was supported by grants from the National Science Foundation (IOS-1829176 and IOS-2145811) and the New Mexico Institute of Mining and Technology.

---

**Figure S1.**
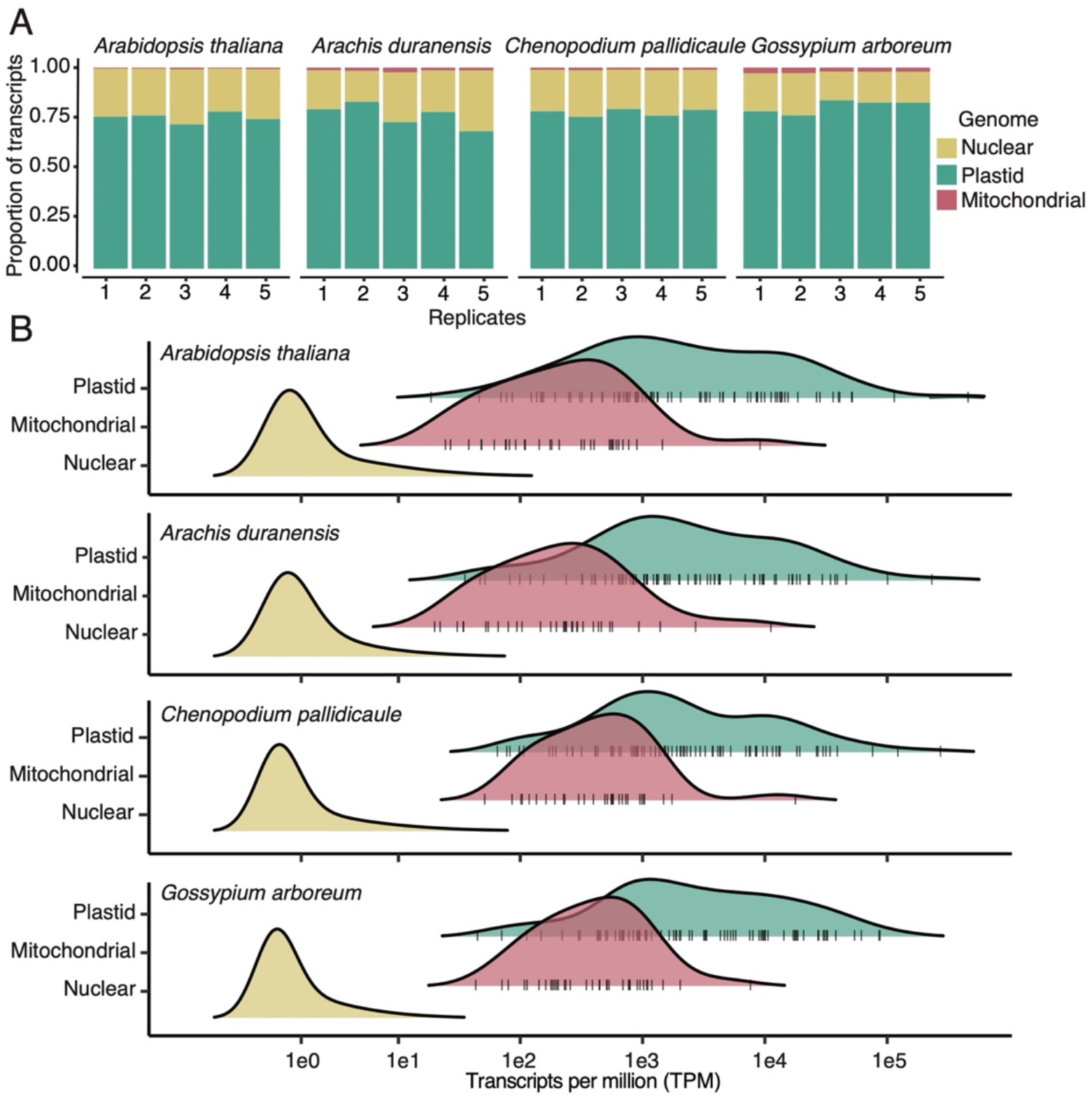
A) Proportion of mRNA transcript by genomic compartment for each biological replicate in four diploid species that represent models of the maternal progenitors of the four allopolyploids shown in Fig. 2. B) Frequency distributions for transcript abundances by gene for each genomic compartment in each species. Vertical tick marks underlying the plastid and mitochondrial distributions indicate individual gene values. Note that there is a downward bias in the distribution of the nuclear genes because mapping was performed against the polyploid reference, so reads from single diploid genes can be split across two homoeologous genes in the reference. This issue does not affect the polyploid analysis (Fig. 2).

**Figure S2.**
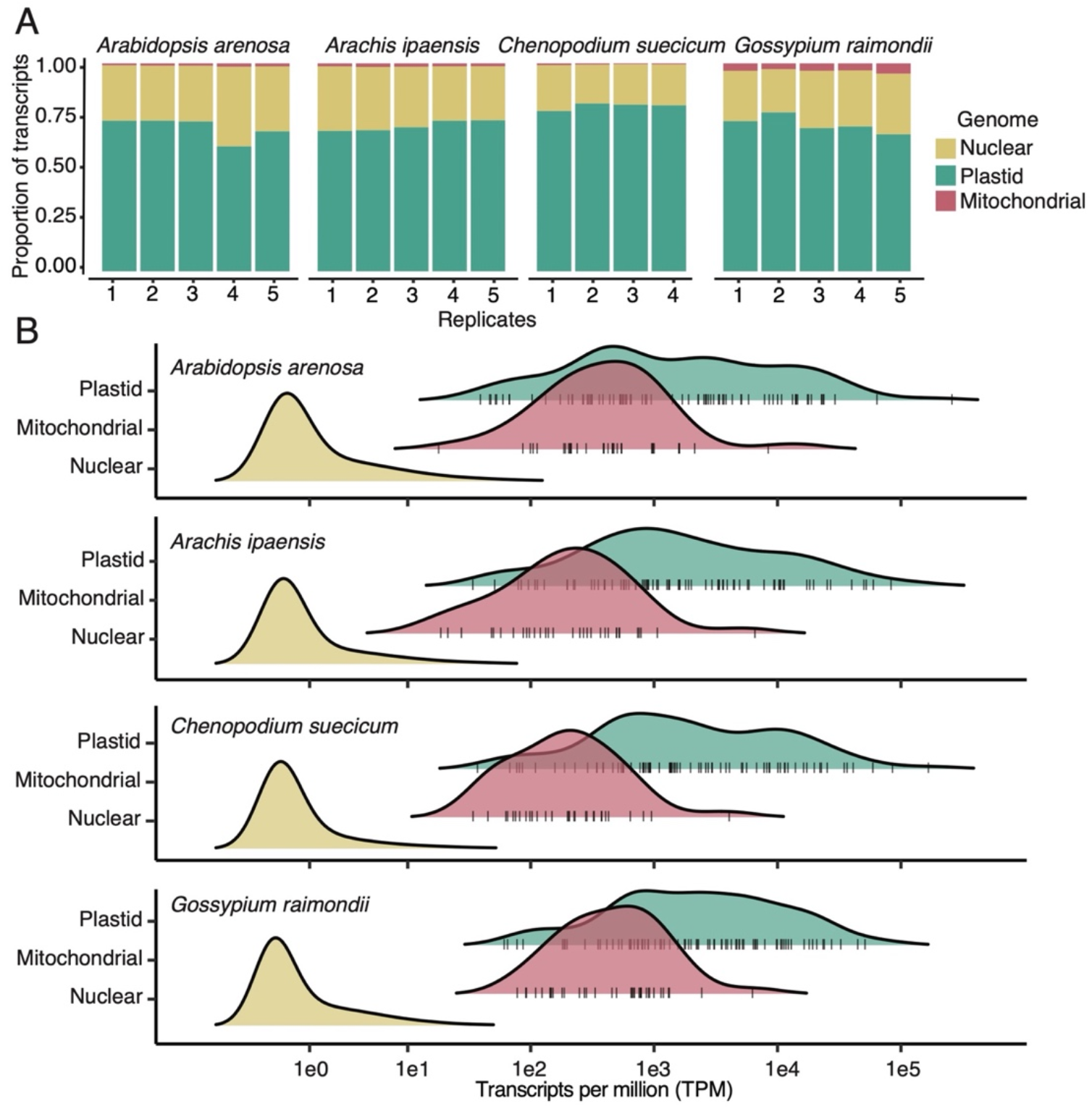
A) Proportion of mRNA transcript by genomic compartment for each biological replicate in four diploid species that represent models of the paternal progenitors of the four allopolyploids shown in Fig. 2. B) Frequency distributions for transcript abundances by gene for each genomic compartment in each species. Vertical tick marks underlying the plastid and mitochondrial distributions indicate individual gene values. Note that there is a downward bias in the distribution of the nuclear genes because mapping was performed against the polyploid reference, so reads from single diploid genes can be split across two homoeologous genes in the reference. This issue does not affect the polyploid analysis (Fig. 2).

**Figure S3.**
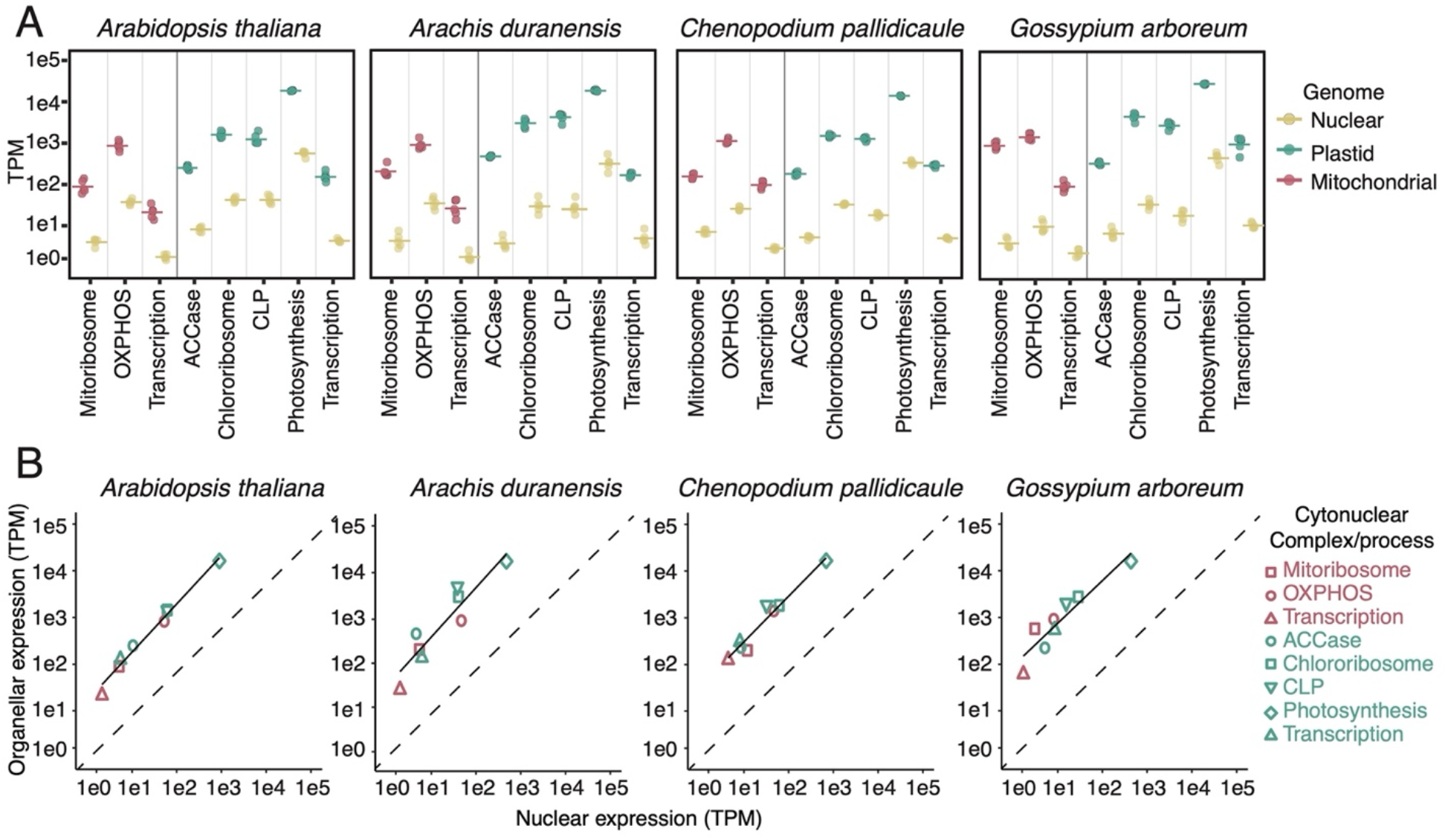
A) Transcript abundance for organellar genes and their interacting N-mt/N-pt counterparts broken down by functional categories according the CyMIRA classification scheme (8) in four diploid species that represent models of the maternal progenitors of the four allopolyploids shown in Fig. 3. Reported values are averaged across genes within a category. Points represent biological replicates, and horizontal lines indicate the mean across replicates. In all cases, organellar mRNA abundance (measured as transcripts per million or TPM) exceeds the corresponding nuclear mRNA abundance by at least ten-fold. B) Correlation between organellar and nuclear mRNA abundances across CyMIRA categories, with the solid line representing a best fit linear model, and the dashed line representing a 1:1 line. Despite the large imbalance towards organellar mRNAs, the cytonuclear ratio remains relatively consistent across functional categories expressed at very different levels, resulting in a strong positive correlation (*p* < 0.001 in all cases). For both sets of plots, the analysis of nuclear genes was limited to those found in homologous “quartets” (see Materials and Methods). Because the reads from these diploids were mapped against a polyploid reference, expression values for the two homoeologs in the polyploid genome were summed for this analysis to avoid loss of reads due to cross-mapping to the other subgenome.

**Figure S4.**
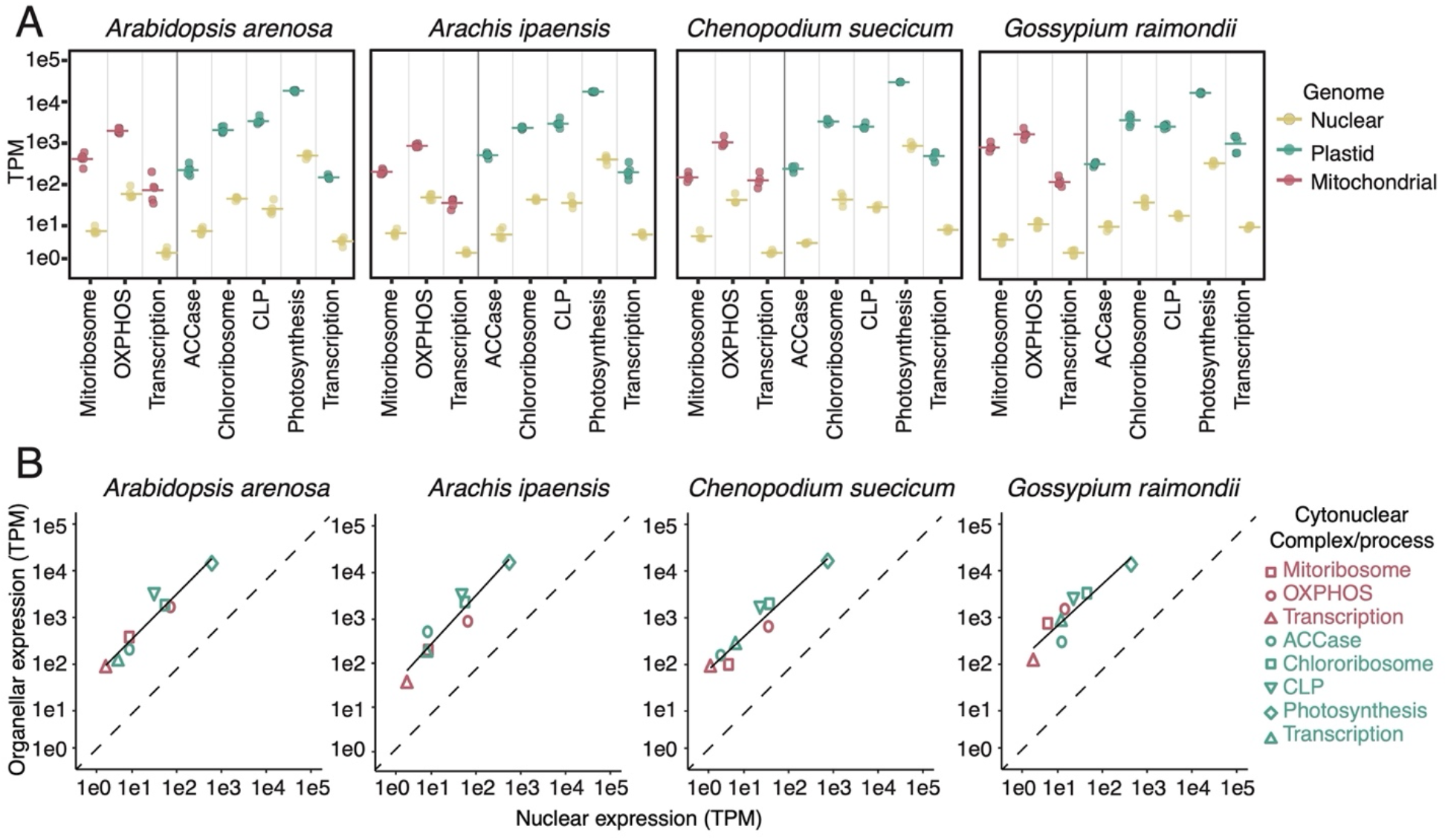
A) Transcript abundance for organellar genes and their interacting N-mt/N-pt counterparts broken down by functional categories according the CyMIRA classification scheme (8) in four diploid species that represent models of the paternal progenitors of the four allopolyploids shown in Fig. 3. Reported values are averaged across genes within a category. Points represent biological replicates, and horizontal lines indicate the mean across replicates. In all cases, organellar mRNA abundance (measured as transcripts per million or TPM) exceeds the corresponding nuclear mRNA abundance by at least ten-fold. B) Correlation between organellar and nuclear mRNA abundances across CyMIRA categories, with the solid line representing a best fit linear model, and the dashed line representing a 1:1 line. Despite the large imbalance towards organellar mRNAs, the cytonuclear ratio remains relatively consistent across functional categories expressed at very different levels, resulting in a strong positive correlation (*p* ≤ 0.001 in all cases). For both sets of plots, the analysis of nuclear genes was limited to those found in homologous “quartets” (see Materials and Methods). Because the reads from these diploids were mapped against a polyploid reference, expression values for the two homoeologs in the polyploid genome were summed for this analysis to avoid loss of reads due to cross-mapping to the other subgenome.

**Figure S5.**
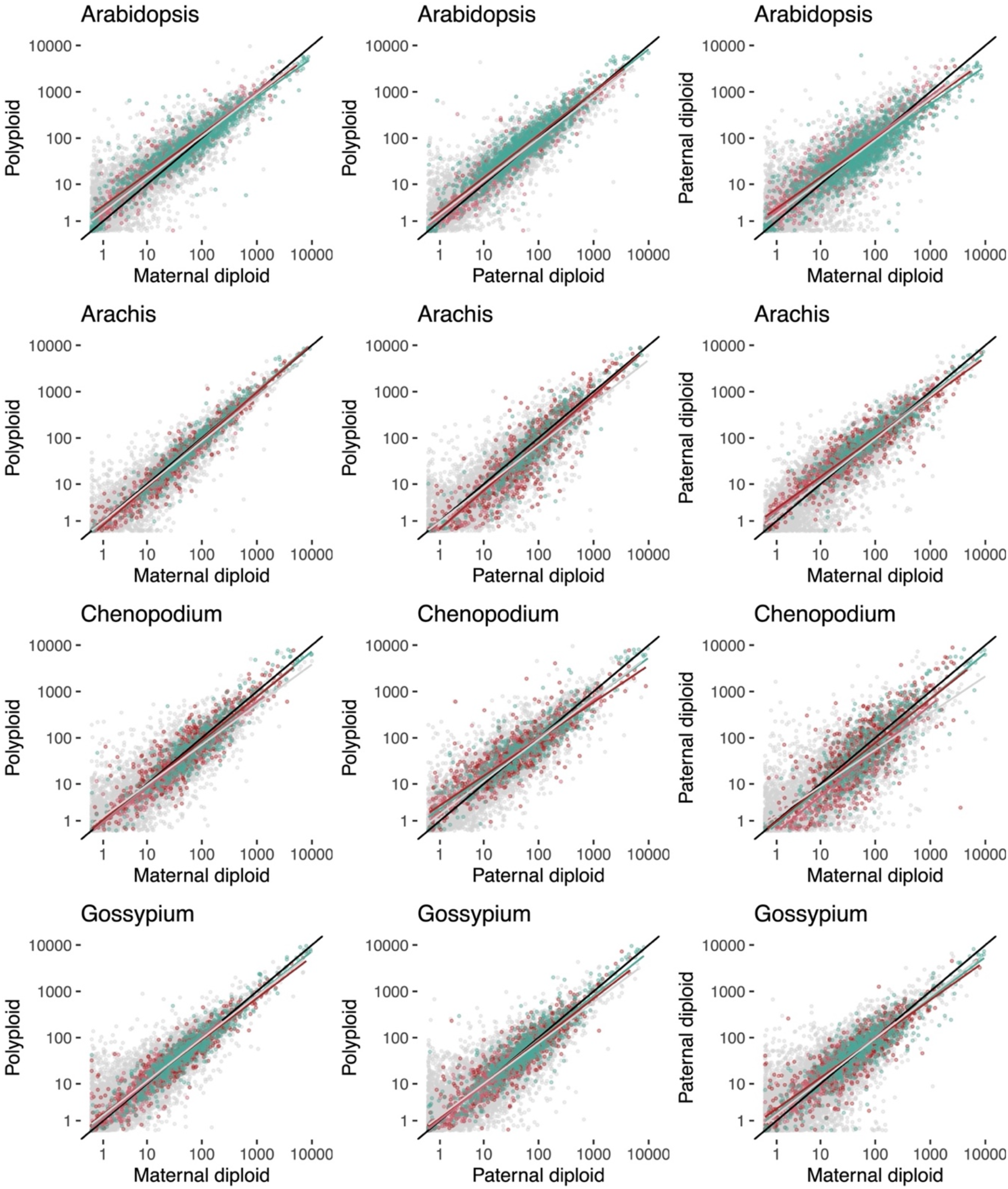
Comparisons between congeneric species in relative abundance of mRNAs, as measured by transcripts per million (TPM). Each point represents an average across biological replicates, with gray, red and green indicating nuclear, mitochondrial, and plastid genes respectively. The black line is the 1:1 line. Colored lines represent best-fit linear models for each gene set separately. Each row corresponds to a different genus. Plots in the left column compare the polyploid against the maternal diploid. Plots in the center column compare the polyploid against the paternal diploid. Plots in the right column compare the paternal diploid against the maternal diploid.

**Table S1.**
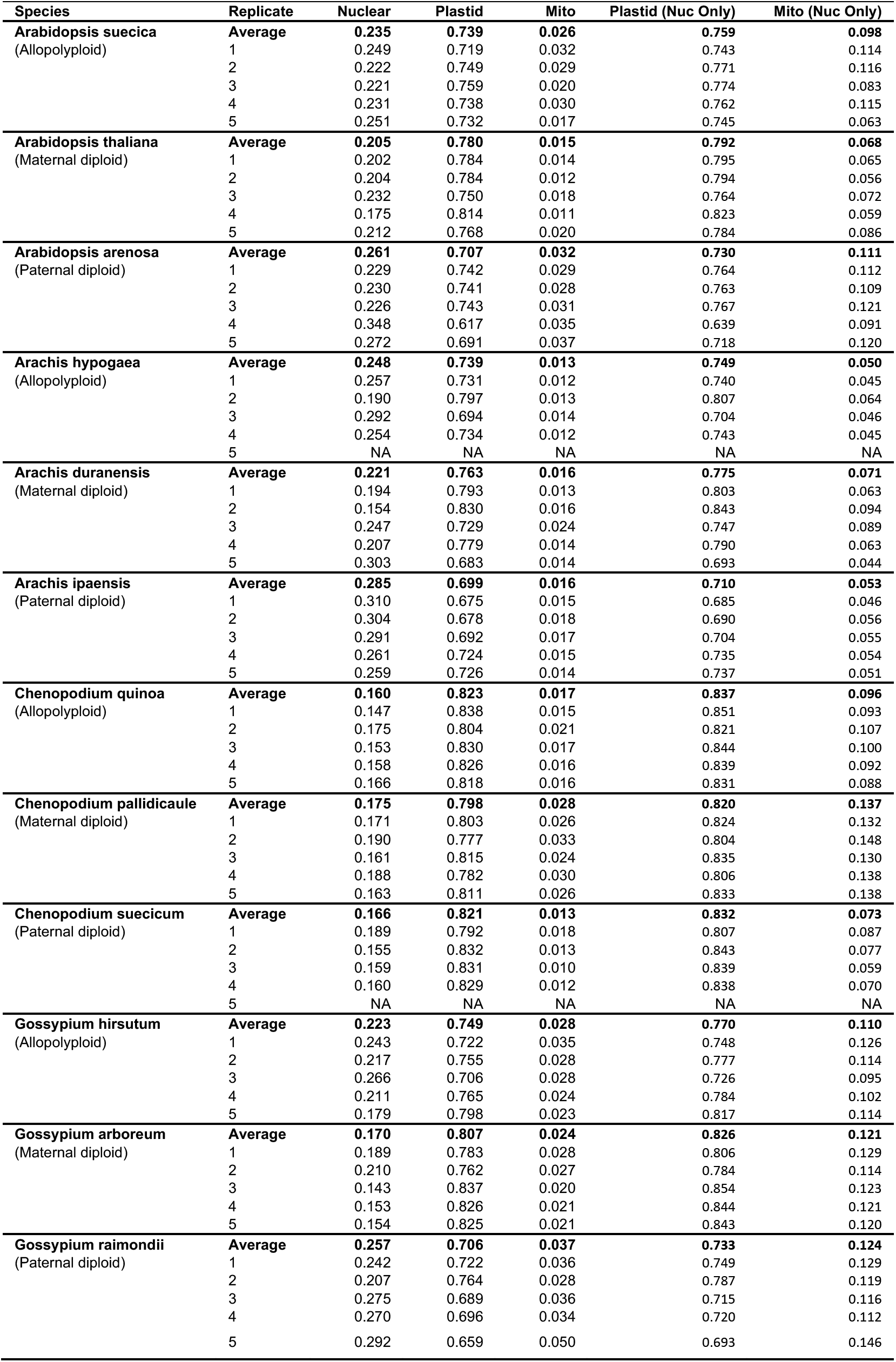
Proportion of mRNAs by genomic compartment. The “Nuc Only” values are calculated for mitochondria and plastids after excluding mRNAs from the other organellar genome, so they only reflect abundance relative to the nucleus.

**Table S2.**
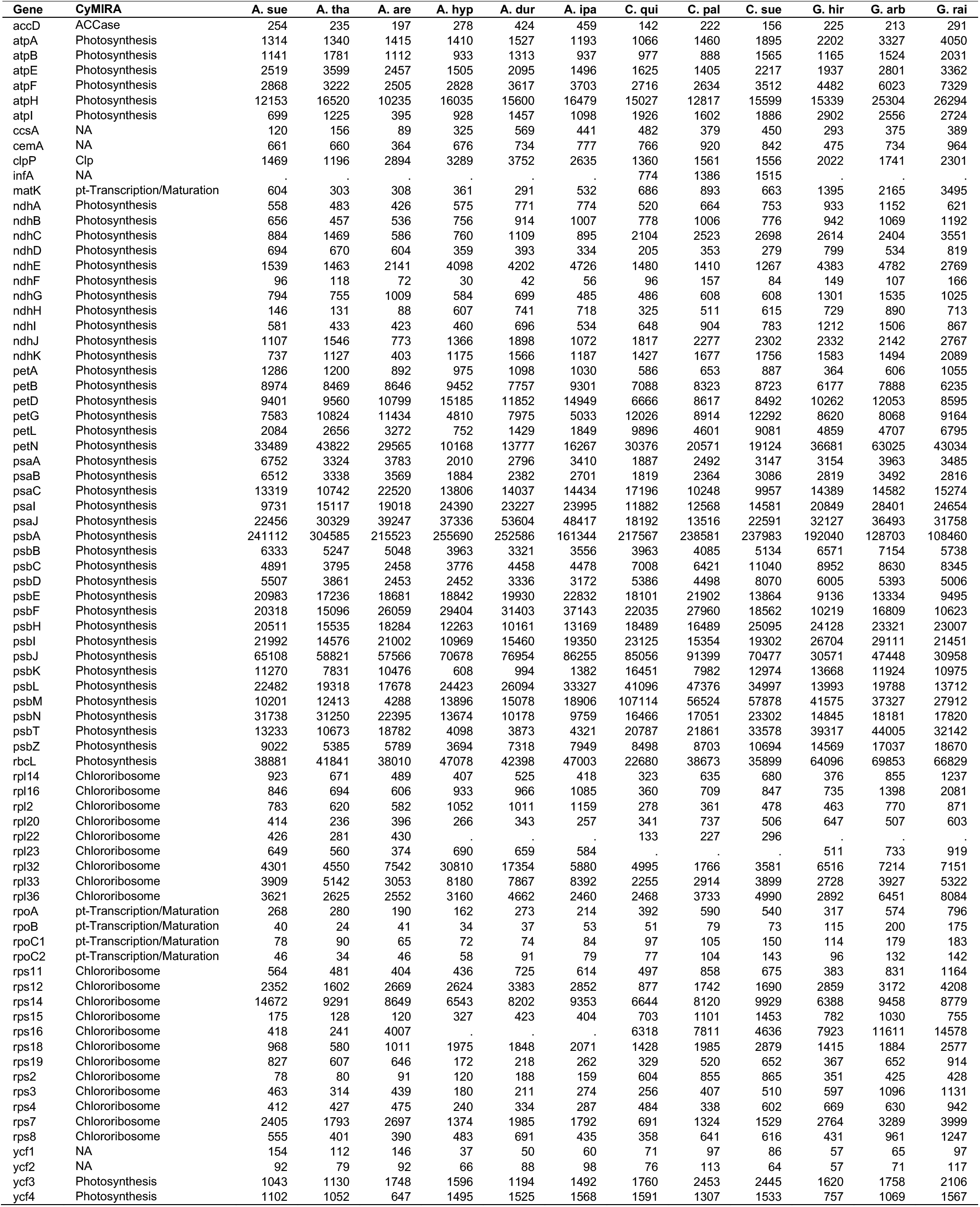
Average transcripts per million (TPM) values by species for each plastid gene

**Table S3.** Average transcripts per million (TPM) values by species for each mitochondrial gene

| Gene | CyMIRA | A. sue | A. tha | A. are | A. hyp | A. dur | A. ipa | C. qui | C. pal | C. sue | G. hir | G. arb | G. rai |
| --- | --- | --- | --- | --- | --- | --- | --- | --- | --- | --- | --- | --- | --- |
| atp1 | OXPHOS | 1769 | 829 | 1104 | 470 | 438 | 489 | 748 | 1123 | 859 | 1390 | 1072 | 2362 |
| atp4 | OXPHOS | 381 | 267 | 411 | 613 | 784 | 737 | 544 | 834 | 447 | 890 | 818 | 1111 |
| atp6 | OXPHOS | 617 | 347 | 362 | 316 | 386 | 290 | 267 | 550 | 305 | 825 | 858 | 1127 |
| atp8 | OXPHOS | 1162 | 675 | 2030 | 251 | 231 | 331 | 850 | 1010 | 817 | 450 | 388 | 673 |
| atp9 | OXPHOS | 11853 | 6847 | 15219 | 4551 | 6582 | 6099 | 8755 | 12567 | 4627 | 8816 | 5273 | 9081 |
| ccmB | NA | 56 | 47 | 92 | 18 | 49 | 40 | 86 | 82 | 60 | 108 | 97 | 188 |
| ccmC | NA | 134 | 57 | 217 | 175 | 44 | 107 | 242 | 423 | 246 | 231 | 195 | 341 |
| ccmFc | NA | 314 | 74 | 191 | 49 | 28 | 65 | 13 | 144 | 66 | 116 | 74 | 102 |
| ccmFn | NA | . | . | . | 17 | 81 | 106 | 58 | 125 | 58 | 110 | 135 | 166 |
| ccmFn1 | NA | 38 | 19 | 27 | . | . | . | . | . | . | . | . | . |
| ccmFn2 | NA | 78 | 60 | 370 | . | . | . | . | . | . | . | . | . |
| cob | OXPHOS | 517 | 354 | 711 | 566 | 450 | 562 | 114 | 511 | 243 | 251 | 355 | 461 |
| cox1 | OXPHOS | 1746 | 999 | 1217 | 376 | 248 | 268 | 131 | 400 | 355 | 1098 | 1078 | 1505 |
| cox2 | OXPHOS | 486 | 534 | 833 | 567 | 684 | 746 | 799 | 1080 | 668 | 557 | 612 | 842 |
| cox3 | OXPHOS | 609 | 380 | 1362 | 881 | 878 | 1034 | 461 | 1099 | 630 | 878 | 1107 | 1093 |
| matR | mt-Transcription/Maturation | 64 | 21 | 78 | 25 | 26 | 34 | 29 | 122 | 85 | 98 | 60 | 110 |
| mttB | NA | 122 | 33 | 181 | 52 | 36 | 18 | 96 | 149 | 58 | 470 | 172 | 263 |
| nad1 | OXPHOS | 458 | 276 | 657 | 184 | 191 | 234 | 627 | 1100 | 404 | 610 | 416 | 763 |
| nad2 | OXPHOS | 246 | 146 | 311 | 106 | 94 | 142 | 298 | 699 | 196 | 304 | 223 | 482 |
| nad3 | OXPHOS | 542 | 311 | 179 | 61 | 97 | 110 | 135 | 121 | 158 | 273 | 251 | 396 |
| nad4 | OXPHOS | 244 | 222 | 446 | 240 | 203 | 379 | 162 | 358 | 152 | 401 | 386 | 826 |
| nad4L | OXPHOS | 487 | 596 | 969 | 321 | 536 | 657 | 232 | 384 | 299 | 920 | 570 | 1667 |
| nad5 | OXPHOS | . | . | . | 197 | 155 | 189 | 108 | 465 | 204 | 464 | 345 | 551 |
| nad6 | OXPHOS | 674 | 607 | 969 | 290 | 246 | 248 | 320 | 819 | 269 | 702 | 963 | 1721 |
| nad7 | OXPHOS | 659 | 225 | 696 | 236 | 403 | 480 | 274 | 1049 | 716 | 574 | 630 | 982 |
| nad9 | OXPHOS | 1056 | 406 | 945 | 835 | 2099 | 942 | 641 | 1004 | 545 | 955 | 992 | 1728 |
| rpl10 | Mitoribosome | . | . | . | . | . | . | . | . | . | 852 | 758 | 1070 |
| rpl16 | Mitoribosome | 210 | 135 | 721 | 101 | 133 | 54 | . | . | . | 603 | 615 | 459 |
| rpl2 | Mitoribosome | 539 | 119 | 616 | . | . | . | . | . | . | 193 | 124 | 302 |
| rpl5 | Mitoribosome | 75 | 109 | 226 | 160 | 99 | 159 | 258 | 402 | 212 | 204 | 146 | 329 |
| rps1 | Mitoribosome | . | . | . | 79 | 71 | 110 | . | . | . | . | . | . |
| rps10 | Mitoribosome | . | . | . | 372 | 217 | 358 | . | . | . | 1953 | 2102 | 2726 |
| rps12 | Mitoribosome | 151 | 120 | 203 | 169 | 262 | 287 | 191 | 177 | 182 | 347 | 376 | 445 |
| rps13 | Mitoribosome | . | . | . | . | . | . | 242 | 294 | 190 | . | . | . |
| rps14 | Mitoribosome | . | . | . | 67 | 74 | 152 | . | . | . | 583 | 622 | 933 |
| rps3 | Mitoribosome | 160 | 83 | 368 | 22 | 29 | 23 | 78 | 98 | 59 | 283 | 229 | 253 |
| rps4 | Mitoribosome | 92 | 37 | 86 | 218 | 437 | 230 | 73 | 251 | 60 | 195 | 229 | 196 |
| rps7 | Mitoribosome | 52 | 25 | 212 | . | . | . | 106 | 248 | 91 | 112 | 109 | 244 |
| sdh4 | OXPHOS | . | . | . | . | . | . | . | . | . | 921 | 1207 | 1218 |

**Table S4.** *R*^2^ values for correlations between species pairs for log TPM values from organellar genes (see Fig. 4).

| **Genus** | **Genome** | **Polyploid vs.**<br>**Maternal Diploid** | **Polyploid vs.**<br>**Paternal Diploid** | **Maternal Diploid**<br>**vs. Paternal Diploid** |
| --- | --- | --- | --- | --- |
| Arabidopsis | Plastid | 0.976 | 0.948 | 0.927 |
| Arabidopsis | Mito | 0.885 | 0.780 | 0.804 |
| Arachis | Plastid | 0.986 | 0.972 | 0.979 |
| Arachis | Mito | 0.824 | 0.841 | 0.884 |
| Chenopodium | Plastid | 0.970 | 0.972 | 0.973 |
| Chenopodium | Mito | 0.791 | 0.811 | 0.871 |
| Gossypium | Plastid | 0.977 | 0.954 | 0.976 |
| Gossypium | Mito | 0.917 | 0.906 | 0.910 |

**Table S5.**
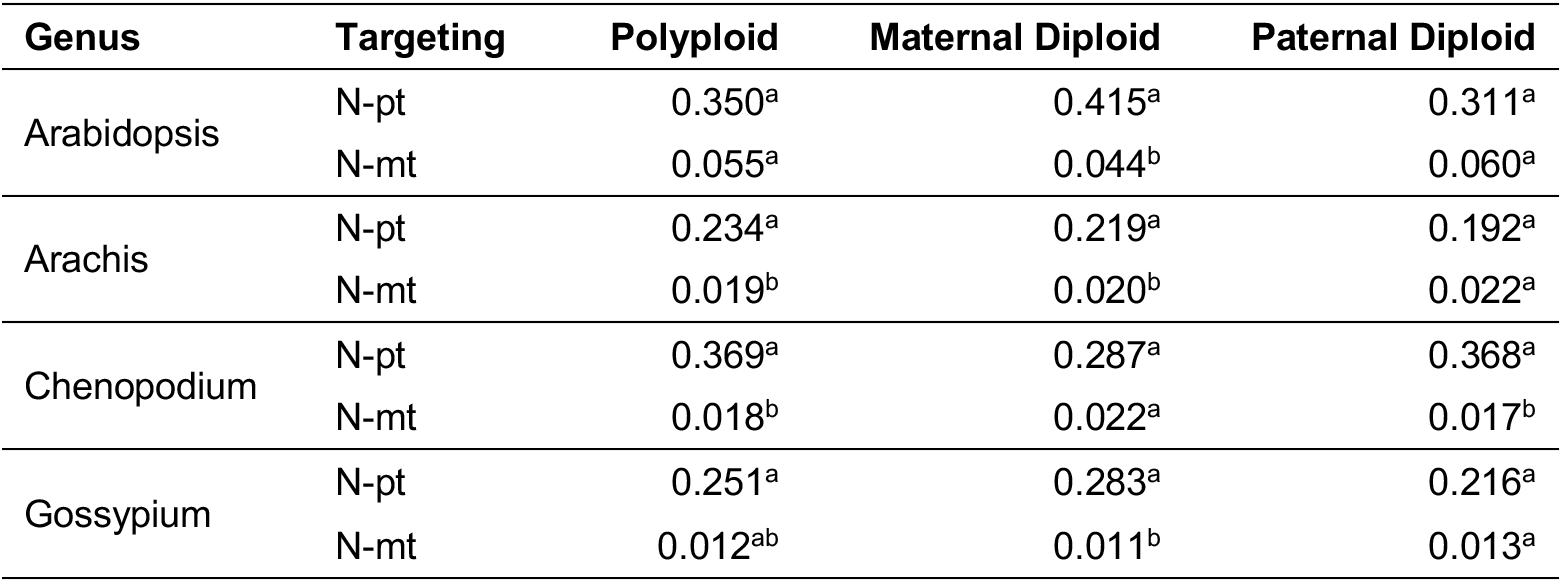
The proportion of nuclear mRNA transcripts derived from N-pt and N-mt genes averaged across biological replicates. The analysis was limited to nuclear genes found in homologous “quartets” (see Materials and Methods). Expression values for the two homoeologs in the polyploid reference genome were summed for this analysis. Superscript letters summarize significant differences between species based on *post hoc* pairwise comparisons (species that share a letter are not significantly different from each other). Models were implemented for each genus and genome separately, so pairwise comparisons should only be interpreted within a row.

**Table S6.** Estimated marginal means by species for log10 transcript per million (TPM) values from mixed-effects models of gene-specific expression levels. Superscript letters summarize significant differences between species based on *post hoc* pairwise comparisons (species that share a letter are not significantly different from each other). Models were implemented for each genus and genome separately, so pairwise comparisons should only be interpreted within a row.

| Genus | Genome | Polyploid | Maternal Diploid | Paternal Diploid |
| --- | --- | --- | --- | --- |
| Arabidopsis | Plastid | 3.26<sup>a</sup> | 3.21<sup>b</sup> | 3.22<sup>b</sup> |
|  | Mito | 3.07<sup>b</sup> | 2.89<sup>c</sup> | 3.15<sup>a</sup> |
| Arachis | Plastid | 3.21<sup>b</sup> | 3.29<sup>a</sup> | 3.28<sup>a</sup> |
|  | Mito | 2.82<sup>c</sup> | 2.90<sup>a</sup> | 2.86<sup>b</sup> |
| Chenopodium | Plastid | 3.27<sup>c</sup> | 3.34<sup>b</sup> | 3.38<sup>a</sup> |
|  | Mito | 3.08<sup>b</sup> | 3.33<sup>a</sup> | 3.10<sup>b</sup> |
| Gossypium | Plastid | 3.37<sup>c</sup> | 3.46<sup>b</sup> | 3.49<sup>a</sup> |
|  | Mito | 3.26<sup>b</sup> | 3.32<sup>a</sup> | 3.33<sup>a</sup> |

